# Pivotal role of PDE10A in the integration of dopamine signals in mice striatal D1 and D2 medium-sized spiny neurones

**DOI:** 10.1101/2021.04.20.440459

**Authors:** Élia Mota, Ségolène Bompierre, Dahdjim Betolngar, Liliana R.V. Castro, Pierre Vincent

## Abstract

Dopamine in the striatum plays a crucial role in reward processes and action selection. Dopamine signals are transduced by D_1_ and D_2_ dopamine receptors which trigger mirror effects through the cAMP/PKA signalling cascade in D1 and D2 medium-sized spiny neurones (MSNs). Phosphodiesterases (PDEs), which determine the profile of cAMP signals, are highly expressed in MSNs, but their respective roles in dopamine signal integration remain poorly understood. We used genetically-encoded FRET biosensors to monitor at the single cell level the functional contribution of PDE2A, PDE4 and PDE10A in the changes of the cAMP/PKA response to transient and continuous dopamine in mouse striatal brain slices. We found that PDE2A, PDE4 and PDE10A operate on the moderate to high cAMP levels elicited by D_1_ or A_2A_ receptor stimulation. In contrast, only PDE10A is able to reduce cAMP down to baseline in both type of neurones, leading to the dephosphorylation of PKA substrates. PDE10A is therefore critically required for dopamine signal integration in both D1 and D2 MSNs.

## Introduction

Dopamine neurotransmission in the striatum is involved in reward-mediated learning and action selection (Surmeier et al., 2007; Kreitzer & Malenka, 2008; Cerovic et al., 2013; Klaus, Alves da Silva, & Costa, 2019). Stimuli predicting a reward trigger the phasic release of dopamine and the timely integration of transient dopamine signal is an important determinant of its action (Gonon, 1997; Day et al., 2007; Schultz, Dayan, & Montague, 1997). Dopamine signals are integrated via the 3’,5’-cyclic adenosine monophosphate (cAMP) - protein kinase A (PKA) signalling pathway, and 3’,5’-cyclic adenosine monophosphate-specific phosphodiesterases (PDEs), which degrade cAMP, are therefore expected to play an important role. Thus, the transient activation of D_1_ receptors triggers an increase in cAMP production, which is terminated by PDE activities. Symmetrically, activation of D_2_ receptors shuts off adenylyl cyclase activity and PDE activities are required to degrade cAMP.

The main target of cAMP in striatal neurones is PKA. The proper transduction of a dopamine signal therefore requires that, after adenylyl cyclase is switched off, cAMP level should be lowered in a concentration range where PKA shows little or no activity. Striatal neurones mainly express PKA regulatory subunits of type IIβ (Cadd & McKnight, 1989), which is activated by cAMP in a 0.1-1 μM concentration range (Dostmann & Taylor, 1991). As highlighted by previous theoretical work (Neves-Zaph, 2017), PDEs with specific Km, Vmax, expression level and subcellular localisation are therefore expected to play different functional roles in the integration of dopamine signals. In this context, we wanted to determine whether the different PDEs expressed in the striatum contributed differently to the integration of dopamine signals.

In situ hybridisation and qPCR reported a particularly robust mRNA expression of the Pde1b, Pde2a, Pde7 and Pde10a genes, and a more marginal expression of Pde4b and Pde8b (Lakics, Karran, & Boess, 2010; Kelly et al., 2014). Pde7b mRNA is highly expressed, but the presence of PDE7B protein has not been reported in the striatum so far. The PDE1B protein is highly expressed in MSNs (Yan et al., 1994; Polli & Kincaid, 1994). PDE1B has a Km of 33 μM for cAMP (Poppe et al., 2008). Our previous work showed that, in D1 MSNs, PDE1 efficiently degrades high cAMP levels in the context of dopamine and NMDA coincidence (Betolngar et al., 2019). In the present work, intracellular calcium was not increased and therefore PDE1B did not contribute to the effects we analysed here.

High levels of PDE2A protein have been reported in the striatum (Stephenson et al., 2009; Stephenson et al., 2012). Its activity is potentiated up to 40 fold by cGMP binding to its GAF-B domain (Martins, Mumby, & Beavo, 1982; Martinez et al., 2002; Jäger et al., 2010). PDE2A can thus mediate a cross-talk between the cGMP and cAMP pathways, as demonstrated in a previous study (Polito et al., 2013). PDE2A has a Km estimated in a 46 - 112 μM range (Sudo et al., 2000; Poppe et al., 2008), suggesting that PDE2A mainly regulates high cAMP levels.

The A, B and D isoforms of PDE4 are also expressed in MSNs (Cherry & Davis, 1999; Perez-Torres et al., 2000; Nishi et al., 2008; Nishi & Snyder, 2010). PDE4B is the main isoform expressed in the striatum, with higher expression in D2 MSNs than in D1 MSNs (Nishi & Snyder, 2010; Nishi et al., 2008). The PDE4 family degrades cAMP with a Km estimated in a 2.5 - 5.5 μM range (Bender & Beavo, 2006; Sudo et al., 2000; Poppe et al., 2008).

The PDE10A protein is present almost exclusively and at very high levels in MSNs (Seeger et al., 2003; Coskran et al., 2006; Lakics et al., 2010), suggesting an important functional role. In contrast to PDE2A and PDE4, PDE10A presents a much lower Km for cAMP, in a range of 0.05-0.25 μM (Wang et al., 2007; Poppe et al., 2008). This suggests a major action of PDE10A on low cAMP levels. Previous studies showed differences between D1 and D2 MSNs in their responsiveness to PDE10A inhibitors, both in vivo and in brain slices (Threlfell et al., 2009; Polito et al., 2015). This D_2_/D_1_ imbalance was consistent with PDE10A inhibitors mimicking the D2 antagonistic action of antipsychotic agents. PDE10A inhibition has therefore been largely studied as a therapeutic strategy in the treatment of schizophrenia or Huntington’s disease (Kehler & Nielsen, 2011; Kehler, Ritzén, & Greve, 2007; Menniti et al., 2007; Chappie et al., 2009; Harada et al., 2020; Schülke & Brandon, 2017). Clinical results however did not confirm these expectations.

In this work, we wanted to delineate the respective functional roles played by PDE2A, PDE4 and PDE10A in the integration of dopamine signals. We used pharmacological inhibitors to selectively block these PDEs while monitoring the cAMP or PKA responses to dopamine with real-time measurement with FRET biosensors. Our data revealed the prominent role played by PDE10A in the integration of dopamine signals through both D_1_ and D_2_ receptors.

## Methods

### Biosensor imaging in brain slice preparations

Mice (male and female C57BL/6J; Janvier labs) were housed under standardised conditions with a 12 hours light/dark cycle, stable temperature (22 ± 1°C), controlled humidity (55 ± 10%) and food and water ad libitum. All animal procedures were performed in accordance with the Sorbonne University animal care committee regulations which complies with the commonly-accepted “3Rs”. Brain slices were prepared from mice aged from 7 to 11 days. Mice were decapitated and the brain quickly removed and immersed in ice-cold solution of the following composition: 125 mM NaCl, 0.4 mM CaCl_2_, 1 mM MgCl_2_, 1.25 mM NaH_2_PO_4_, 26 mM NaHCO_3_, 20 mM glucose, 2.5 mM KCl, 5 mM sodium pyruvate and 1 mM kynurenic acid, saturated with 5% CO_2_ and 95% O_2_. Coronal striatal brain slices were cut with a VT1200S microtome (Leica). The slices were incubated in this solution for 30 minutes and then placed on a Millicell-CM membrane (Millipore) in culture medium (50% Minimum Essential Medium, 50% Hanks’ Balanced Salt Solution, 5.5 g/L glucose, penicillin-streptomycin, Invitrogen). The cAMP biosensor Epac-S^H150^ (Polito et al., 2013) or AKAR4 reporter of PKA/phosphatase equilibrium (Depry, Allen, & Zhang, 2011) were expressed using the Sindbis virus as vector (Ehrengruber et al., 1999): the viral vector was added on the brain slices (~5 x 105 particles per slice), and the infected slices were incubated overnight at 35°C under an atmosphere containing 5% CO_2_. Before the experiment, slices were incubated for 30 min in the recording solution (125 mM NaCl, 2 mM CaCl_2_, 1 mM MgCl_2_, 1.25 mM NaH_2_PO_4_, 26 mM NaHCO_3_, 20 mM glucose, 2.5 mM KCl and 5 mM sodium pyruvate saturated with 5% CO_2_ and 95% O_2_). Recordings were performed with a continuous perfusion of the same solution at 32°C.

Wide-field images were obtained with an Olympus BX50WI or BX51WI upright microscope with a 40x 0.8 NA water-immersion objective and an ORCA-AG camera (Hamamatsu). Images were acquired with iVision (Biovision, Exton, PA, USA). The excitation and dichroic filters were D436/20 and 455dcxt. Signals were acquired by alternating the emission filters, HQ480/40 for donor emission, and D535/40 for acceptor emission, with a filter wheel (Sutter Instruments, Novato, CA, USA). Filters were obtained from Chroma Technology and Semrock.

Photo-release of caged compounds was performed using high power 360 nm LED sources mounted on the epifluorescence port of the microscope, providing 14 mW at the exit of the microscope objective (0.5 s flash duration) or 7.5 mW (1 s flash duration). The combination of LED sources at 360 nm (for uncaging) and 420 nm (off-peak light source for 436 nm excitation filter for biosensor imaging) was purchased from Mightex (Toronto, Canada). The frequency of data acquisition, usually 1 image pair every 30 or 60 seconds, was increased to 0.2 Hz in order to resolve peak dynamics, starting 10 data points before drug uncaging.

### Image analysis

Images were analysed with custom routines written in the IGOR Pro environment (Wavemetrics, Lake Oswego, OR, USA). No correction for bleed-through or direct excitation of the acceptor was applied to keep the benefit of ratiometric calculation, which cancels various imaging artefacts. Biosensor activation level was quantified by ratiometric imaging: donor fluorescence divided by acceptor fluorescence for Epac-S^H150^, and acceptor divided by donor for AKAR4. The emission ratio was calculated for each pixel and displayed in pseudo-colour images, with the ratio value coded in hue and the fluorescence of the preparation coded in intensity. Wide-field imaging allowed for the separation of D1 and D2 MSNs, provided that the infection level was kept low and no fluorescence overlap between neighbouring neurones was observed. Cells were also excluded from the analysis when basal ratio was elevated, when the response to forskolin was lacking or when the neuronal morphology was altered (uneven cell contours). Rare cholinergic interneurons, recognised by their morphology and cell body size larger than 13 μm, were excluded from our analysis.

### Drugs

Solutions were prepared from powders purchased from Sigma-Aldrich (St Quentin-Fallavier, Isère, France). Tetrodotoxin was from Latoxan (Valence, France). BAY607550 was obtained from CliniSciences (Nanterre, France). Other drugs were from Tocris Bio-Techne (Lille, France).

NPEC-DA, (N)-1-(2-Nitrophenyl)ethylcarboxy-3,4-dihydroxyphenethylamine, a chemical precursor of dopamine which can be released by a flash of UV light, was stored at 100 mM concentration in DMSO at −20°C. A new aliquot was used for every day of experiment, kept at 4°C in the dark. NPEC-DA (3 μM) had no effect by itself on the cAMP level in D1 and D2 MSNs. After a few minutes to allow for the diffusion of the compound into the brain slice, a flash of UV light was applied through the objective to convert all NPEC-DA in the imaging field of view into free dopamine. Dopamine concentration in the brain slice then decays exponentially with a time-constant of ~90 s (Yapo et al., 2017).

### Data and statistical analysis

The data and statistical analysis comply with the recommendations on experimental design and analysis in pharmacology (Curtis et al., 2018). Statistical tests were performed using Igor Pro. Responses obtained from all neurones of the same type in one experiment were averaged together and statistics were calculated per experiment (N). (n) indicates the total number of neurones per condition. (A) indicates the number of animals used for preparing brain slices. The alpha level for statistical significance was fixed at 0.05, indicated by * in figures and tables. n.s. indicates that the test was not significant. For some conditions, the Shapiro-Wilk test rejected the hypothesis of normal data distribution and therefore non-parametric statistics were used throughout this study. The Wilcoxon signed rank test was used for paired data. The Wilcoxon-Mann-Whitney test was used to test whether the treatment increased the ratio value for unpaired data. A Kruskal-Wallis followed by Dunn tests were used for more than 2 conditions (Figure 6).

### Estimates of cAMP concentration

Ratiometric imaging can be used to determine absolute analyte concentrations, with ratio values following a Hill equation from Rmin, the ratio in the absence of ligand, to Rmax, the ratio in the presence of saturating ligand (Grynkiewicz, Poenie, & Tsien, 1985). For Epac-S^H150^, ratio values were translated into corresponding cAMP concentration estimates using a Hill equation with a Kd of 4.4 μM and a Hill coefficient of 0.77 (Polito et al., 2013). Rmax was determined for each neurone by the final application of forskolin and IBMX. The basal ratio was estimated by inhibiting adenylyl cyclases with 200 μM SQ22536, which decreased the ratio by 0.037 ratio units to reach the minimal ratio, Rmin. This value was not statistically different between D1 and D2 MSNs (N=8, n=58, A=7). This basal ratio value of 0.037, corresponding to ~50 nM cAMP, was subtracted from baseline ratio when estimating cAMP concentration.

## Results

### Imaging the cAMP/PKA signalling pathway in striatal brain slices

The striatum comprises ~90% of medium-sized spiny neurones (MSNs) distributed in two sub-groups, which express dopamine receptors of either D_1_ or D_2_ type (Valjent et al., 2009; Bertran-Gonzalez et al., 2010). These two sets of neurones will be hereafter called D1 MSNs and D2 MSNs, respectively. D2 MSNs also express adenosine A_2A_ receptors, positively coupled to cAMP production (Fink et al., 1992). A strict segregation of receptors in two separate MSN populations can be readily imaged with wide-field microscopy in mouse striatal brain slices expressing genetically-encoded biosensors for the cAMP/PKA signalling pathway (Yapo et al., 2017; Betolngar et al., 2019). Ratiometric quantification provides real-time monitoring of changes in cAMP/PKA signalling in several MSNs at the same time during the application of drugs in the microscope chamber. The identity of D1 and D2 MSNs in the field of view can be unambiguously revealed by its positive biosensor response to receptor activation: SKF-81297 (SKF, 100 nM), a D_1_-receptor selective agonist, and CGS 21680 (CGS, 1 μM), an A_2A_ agonist, respectively (Figure 1A). At the end of every experiment, the adenylyl cyclases activator forskolin (fsk, 13 μM) together with the non-selective PDE inhibitor IBMX (200 μM) are applied to reach the saturating ratio value, (Rmax) for the biosensor. For each neurone, the ratio trace is normalised between the basal ratio, measured before the application of CGS, and Rmax (Figure 1D). Male and female mice pups were used in this study, and no difference related to sex was observed.

**Figure 1.**
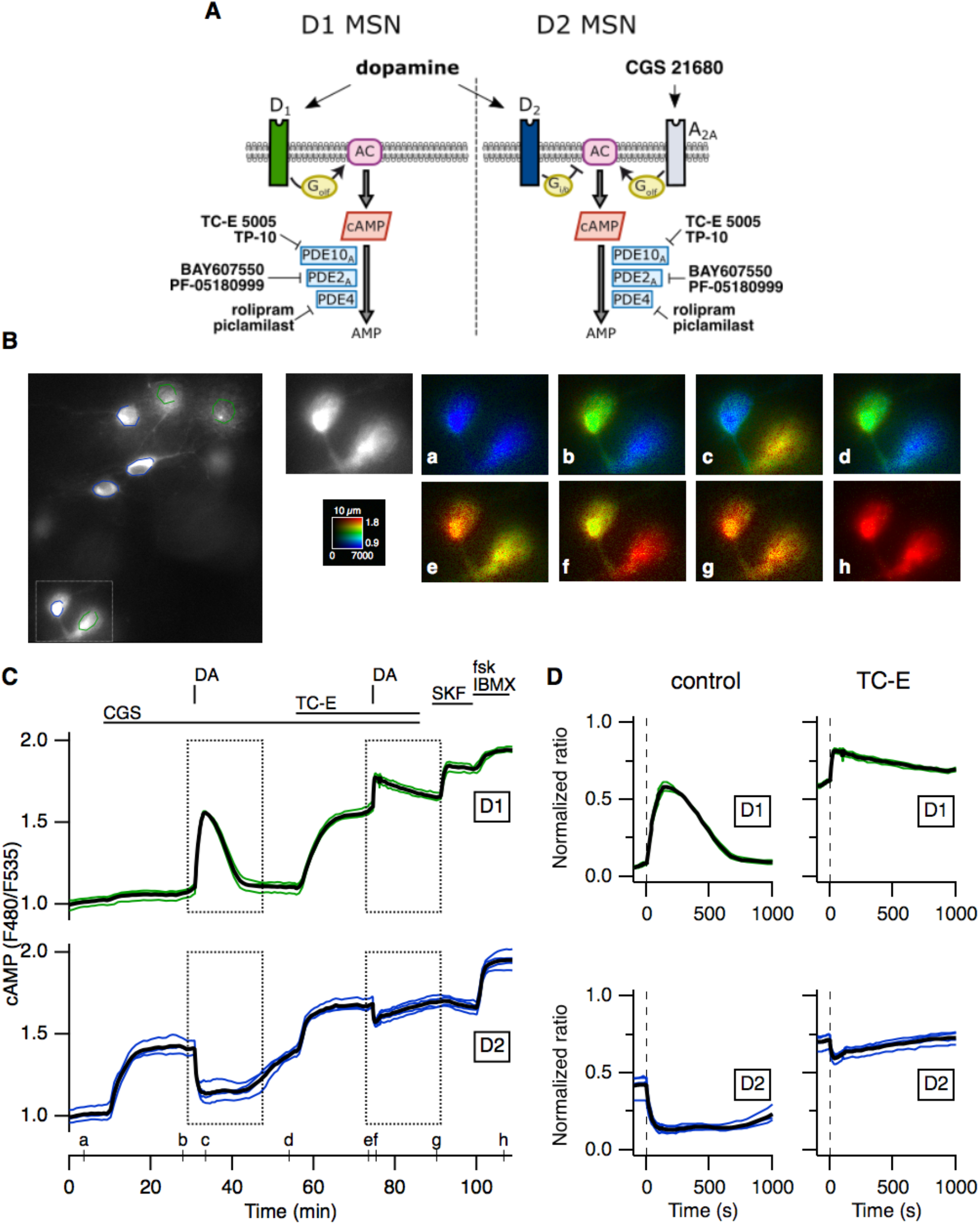
PDE10A activity determines the profile of cAMP responses to dopamine in striatal MSNs. The Epac-S^H150^ biosensor was expressed in mice striatal brain slices and imaged with wide-field microscopy. (A) Schematic representation of cAMP signalling in D1-type and D2-type MSNs. (B) Left: part of the field of view showing the fluorescence at 535 nm (F535). The regions of interest on individual cells define the area used for measuring the ratio (F480/F535). Right: pseudo-colour images (a-h) of the region indicated by the rectangle and corresponding to the time points indicated on the graph in C. (C) Each trace on the graph indicates the emission ratio indicative of intracellular cAMP concentration measured on the cell body of an individual neurone. Drug application is indicated by horizontal bars. The A_2a_ receptor agonist CGS 21680 (CGS, 1 μM) increases cAMP level in D2 MSNs. Dopamine released from caged NPEC-DA (3 *μ*M) by a flash of UV light (DA) triggered transient responses of opposite directions in D1 MSNs (green traces) and D2 MSNs (blue traces). A second dopamine release was applied in the presence of the PDE10A inhibitor, TC-E 5005 (TC-E, 1 *μ*M). Forskolin (fsk, 13 *μ*M) and IBMX (200 *μ*M) maximally increased cAMP levels for each neurone. (D) Traces normalised between baseline and the maximal response to fsk and IBMX; first (ctl, left) and second (TC-E, right) responses to dopamine stimulation. Averages for each type of MSNs are represented as black lines.

### PDE10A degrades low and high cAMP levels

First, we used the Epac-S^H150^ biosensor (Polito et al., 2013) to image the cAMP response to transient dopamine stimulation and analyse PDE10A action on dopamine response (Figure 1B-D). Since dopamine D_2_ receptors are negatively coupled to cAMP production and since the basal level of cAMP is low (see estimate of baseline cAMP level in Methods), intracellular cAMP production was first increased selectively in D2 MSNs by applying the adenosine A_2A_ receptor agonist CGS (Figure 1B a-b, 1C, blue traces on the graph). This mimics the adenosine tone in the striatum. All stimulations with CGS in this study were performed in the presence of the adenosine A_1_ receptor antagonist PSB36 (PSB, 100 nM), and the sodium channel blocker tetrodotoxin (200 nM). Once a steady-state cAMP level was reached in D2 MSNs, a flash of UV light was applied to the field of view to release dopamine (3 μM) from the “caged” precursor NPEC-DA (see details in Methods). This brief dopamine stimulation transiently decreased cAMP level in D2 MSNs (Figure 1B b-c and 1C, blue traces), while it simultaneously increased cAMP in D1 MSNs (Figure 1B b-c and 1C, green traces). These opposite changes in cAMP concentration in D1 and D2 MSNs with this uncaging protocol have already been described, and were shown to be repeatable in the same neurones with the same amplitude and kinetics (Yapo et al., 2017; Betolngar et al., 2019).

To test the functional role of PDE10A on dopamine-induced changes in cAMP levels, a second trial with caged dopamine was performed in the presence of a PDE10A inhibitor. We used TC-E 5005 (TC-E, 1 μM) applied in the bath a few minutes before and maintained during and after the release of dopamine (Figure 1B e-g, 1C). Application of TC-E alone significantly increased the ratio level in D1 MSNs. In D2 MSNs, TC-E potentiated the cAMP response to CGS. On this steady-state cAMP level, the second episode of dopamine uncaging induced a transient cAMP increase in D1 MSNs and a small transient decrease in cAMP in D2 MSNs (Figure 1B e-g). Traces from MSNs of the same type were averaged, and that average normalised between baseline and Rmax (Figure 1D). Time 0 was set at the time of dopamine uncaging.

The same experiment was performed 6 times (N=6), and the average traces from each experiment were averaged and displayed in Figure 2A (green for D1, blue for D2 MSNs). The steady-state level measured before dopamine uncaging at time 0, and the peak response to dopamine, were plotted with connecting lines to show for each experiment the ratio change resulting from TC-E treatment. Normalised ratio values can be converted to an estimated cAMP concentration (see Methods), indicated by a scale on the right of the graph. Numerical values are indicated in Table 1.

**Figure 2:**
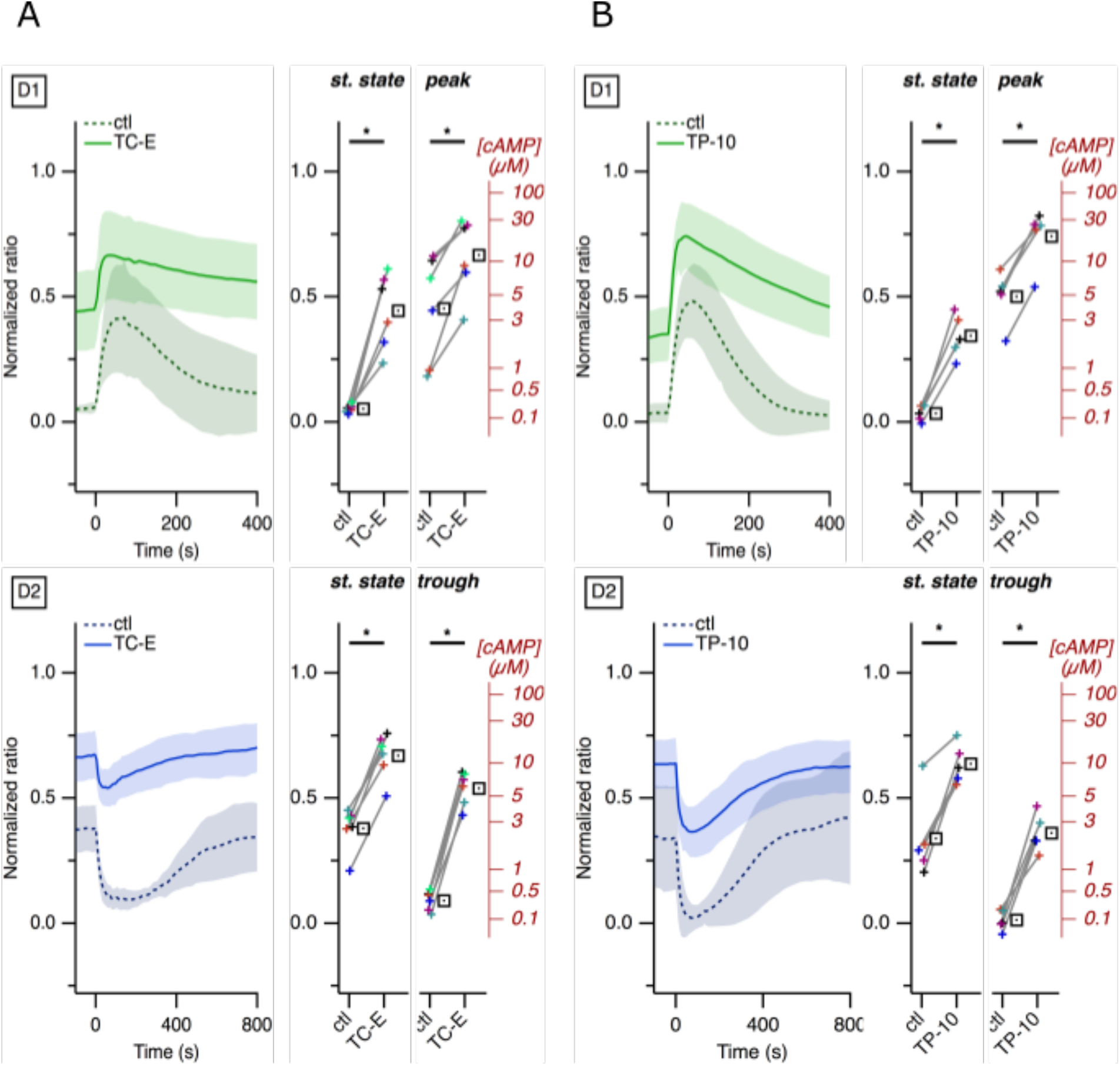
PDE10A activity is required to lower cAMP levels to baseline in D1 and D2 MSNs. Same protocol as in Figure 1. PDE10A was inhibited either with (A) TC-E (1 *μ*M, N=6)) or with (B) TP-10 (1 *μ*M, N=5). Traces show the average ratio response to dopamine in D1 and D2 MSNs, without (ctl, dashed line) or with PDE10A inhibitor (plain line). Shading indicates 95% confidence interval. Plots show the average of the steady-state ratio level before dopamine uncaging (st. state) and the peak (D1) or trough (D2) amplitude of dopamine responses alone (ctl, control) or in the presence of TC-E or TP-10. The average of the different repeats is represented with a square symbol. The axis on the right of the plot indicates the cAMP concentration estimated from the ratio level. * indicates a statistically significant difference.

**Table 1:**
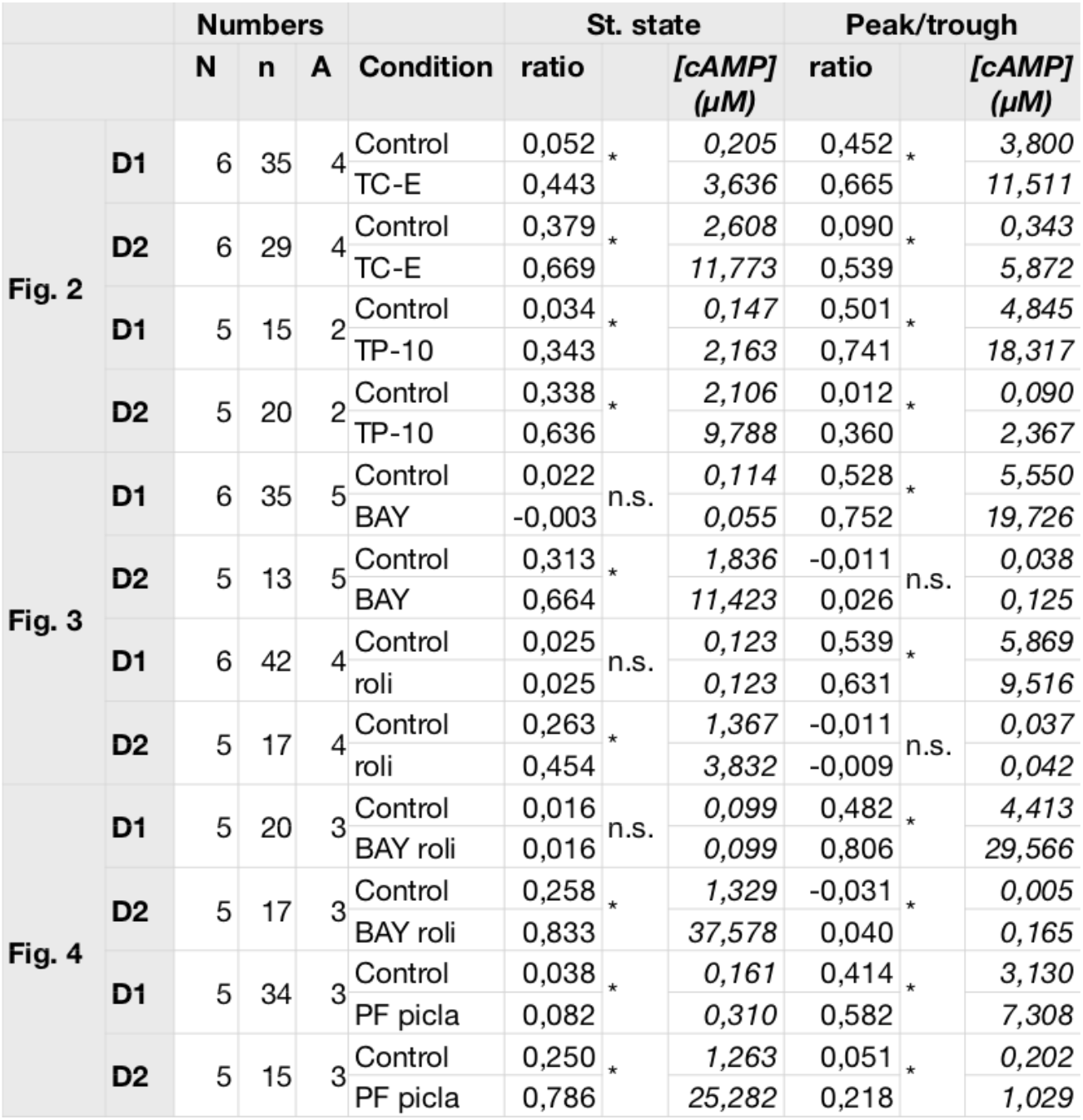
Measurements corresponding to figures 2–4. N indicates the number of experiments; n indicates the total number of cells; A indicates the number of different animals. * indicates a statistically significant difference; n.s. indicates that the difference was not significant.

In D1 MSNs, the steady-state level before dopamine uncaging (Figure 2A, “st. state”) was increased by TC-E treatment, with corresponding cAMP concentration rising from ~0.2 to ~4 μM. TC-E also increased the peak response to dopamine uncaging, from ~4 to ~12 μM.

In D2 MSNs, the application of TC-E increased the steady-state ratio level elicited by A_2A_ receptor activation from ~3 to ~12 μM. The trough in cAMP level induced by dopamine uncaging was reaching ~0.3 μM in control conditions, while in the presence of TC-E, that level was reduced only to ~5 μM, far from reaching the baseline. We also tested the effect of TP-10 (1 μM), another potent and highly selective PDE10A inhibitor and obtained similar results (Figure 2B and Table 1).

These experiments show that PDE10A is an important regulator of basal cAMP level in D1 MSNs, as well as in the regulation of peak cAMP level during a positive cAMP response to dopamine. In D2 MSNs, PDE10A participates in the regulation of the steady-state level reached upon A_2A_ receptor activation. PDE10A activity also appears required to decrease cAMP concentration toward baseline level upon D_2_ receptor activation, with cAMP levels remaining above 2 μM.

### PDE2A and PDE4 contribution

We then evaluated the contribution of PDE2A using the same protocol as in Figure 1, except that, instead of inhibiting PDE10A, we selectively inhibited PDE2A with BAY607550 (BAY, 200 nM, Figure 3A). In contrast to PDE10A inhibition, application of BAY had no effect on basal ratio in D1 MSNs. However, it increased the amplitude of the cAMP response to transient dopamine (Figure 3 C and Table 1).

**Figure 3:**
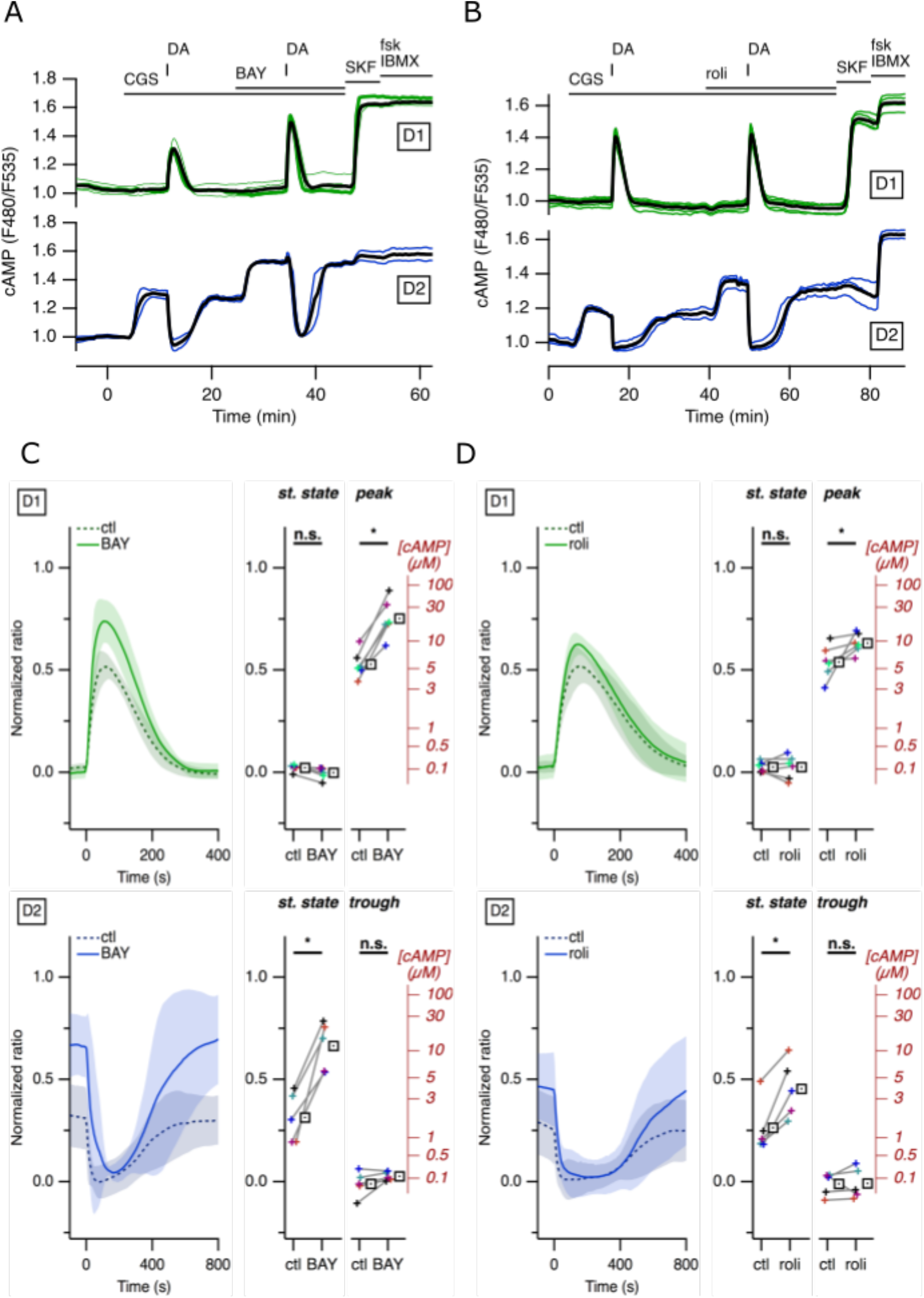
Although PDE2A or PDE4 activities regulate high cAMP levels, they are not required to lower cAMP levels to baseline. Same experiment as in Figure 1, except that PDE2A or PDE4 were inhibited either with (A) BAY607550 (BAY, 200 nM) or with (B) Rolipram (roli, 100 nM), respectively. C and D represent the average of repeats of the same experiments.Statistical tests that showed no difference are indicated with n.s.

In D2 MSNs, the application of BAY on top of the steady-state A_2A_ response increased the ratio. In contrast to PDE10A inhibition, dopamine uncaging in the presence of BAY decreased cAMP level down to baseline (Figure 3 A, C): the minimal ratio level after dopamine release in the presence of BAY was not different from control condition.

To evaluate the contribution of PDE4, the same experiment was repeated using a selective PDE4 inhibitor, Rolipram (roli, 1 μM). The effect of roli was similar to that of BAY (Figure 3 B, D).

These experiments show that PDE2A and PDE4 are both active in the regulation of the peak cAMP response to dopamine in D1 MSNs. These PDEs also contribute to the moderation of the steady-state cAMP response to A_2A_ receptor stimulation in D2 MSNs. However, when either PDE is blocked, D_2_ receptor activation still leads to a decrease of cAMP concentration down to baseline level.

Previous work in cardiomyocytes showed that different phosphodiesterases can act in synergy to regulate cAMP signals (Mika et al., 2019), and PDE2A-PDE4 synergy has been observed in memory processes (Paes et al., 2021). To test for the coincident activity of PDE2A and PDE4, both activities were blocked with PDE2A and PDE4 inhibitors (Figure 4). To rule out any unspecific drug effects, the same protocol was tested with either BAY and roli, or with the PDE2A inhibitor PF-05180999 (PF, 1 μM) and PDE4 inhibitor piclamilast (picla, 1 μM).

**Figure 4:**
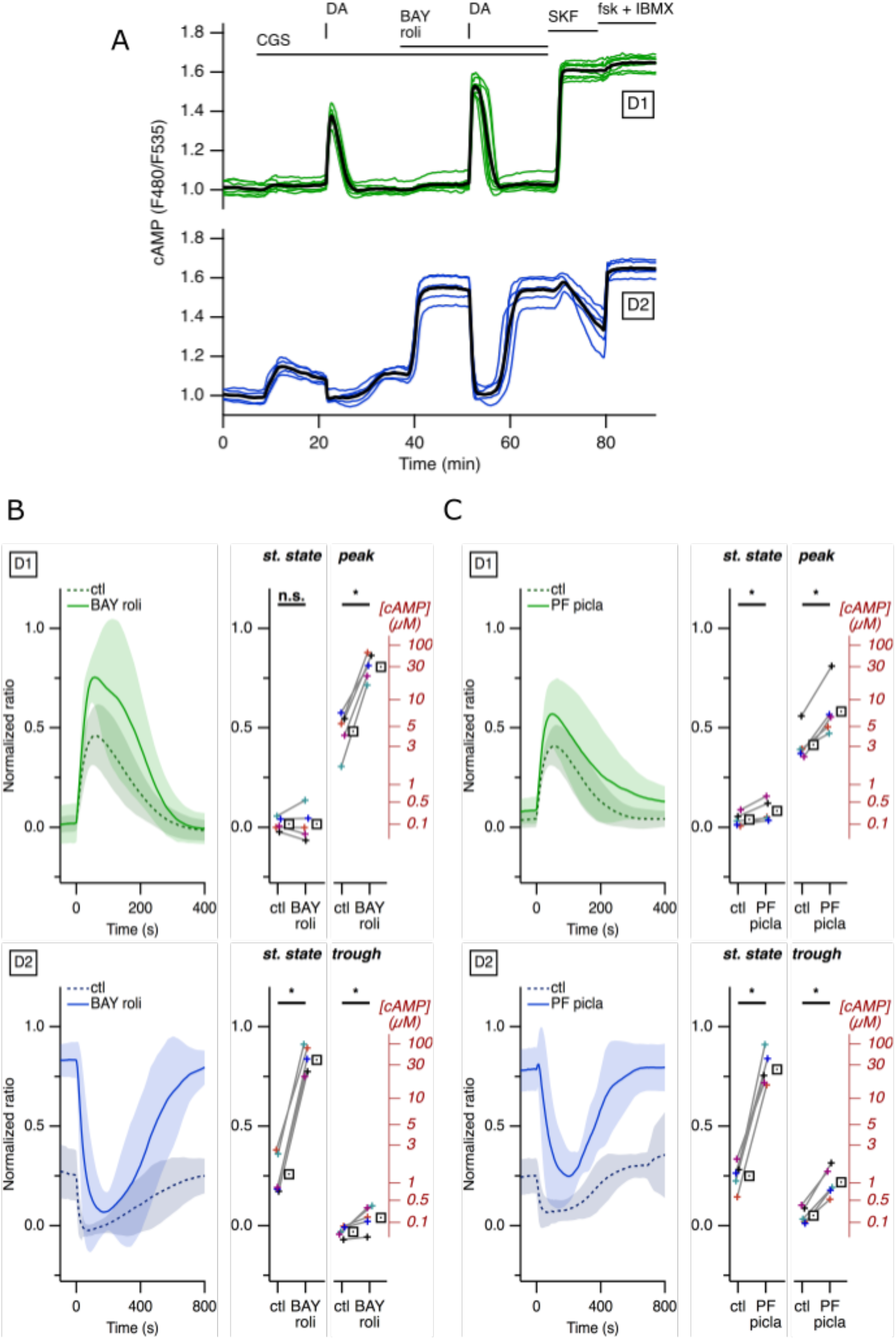
When both PDE2A and PDE4 are inhibited, cAMP can still be degraded to low levels. (A) Same experiment as in Figure 1, except that PDE2A and PDE4 were concomitantly inhibited with BAY607550 (BAY, 200 nM) and Rolipram (roli, 100 nM), respectively. B and C represent the average of repeats of the same experiments, with either (B) BAY and roli, or (C) another combination of selective inhibitors of PDE2 PF-05180999 (PF, 1 *μ*M) and PDE4 piclamilast (picla, 1 *μ*M).

Inhibition of both PDE2A and PDE4 with BAY and roli had no effect on baseline cAMP level in D1 MSNs (Figure 4 A and 4B, green traces), while PF and picla induced a significant, but small, cAMP increase (~0.3 μM, Figure 4C). As expected from the potentiation of the D_1_ response obtained when either PDE was inhibited, the dopamine peak in D1 MSNs was significantly larger when both PDE2A and PDE4 activities were blocked.

Simultaneous inhibition of PDE2A and PDE4 also potentiated the cAMP response to A_2A_ receptor stimulation in D2 MSNs (Figure 4A and 4B, blue traces). When PDE2A and PDE4 were simultaneous inhibited, D_2_ receptor activation still led to a marked decrease in cAMP concentration. With BAY and roli, D_2_ receptor activation decreased cAMP level to a level that was not different from baseline. With PF and picla, cAMP decreased markedly, although remained higher than baseline level, at ~1 μM.

These experiments show that PDE2A and PDE4 act synergistically: their individual activities have minor consequences in the profile of dopamine responses in D1 and D2 MSNs, but they jointly regulate high cAMP levels in D1 and D2 MSNs.

### Integration at the PKA-dependent phosphorylation level

PDE2A and PDE4 thus appear to regulate high cAMP levels reached in D1 MSNs after transient stimulation with dopamine, as well as high cAMP steady-state levels induced by A_2A_ adenosine receptor in D2 MSNs. However, if PDE10A is inhibited, PDE2A and PDE4 activities are not sufficient to lower cAMP concentration below the μM range. cAMP degradation by PDE10A activity therefore appears critical to allow for PKA de-activation and this hypothesis was tested using a PKA-specific biosensor. We monitored the phosphorylation level of PKA substrates using the AKAR4 biosensor, an improved biosensor for monitoring the equilibrium between PKA and phosphatases activities (Depry et al., 2011). PKA activity was first increased in D2 MSNs using CGS in the presence of PSB (Figure 5A a-b), increasing the ratio to the maximal level. Dopamine uncaging (NPEC-DA, 3 μM) produced a transient ratio increase in the D1 MSNs while D2 MSNs displayed a transient ratio decrease (Figure 5). These measurements are consistent with our previous results obtained with the AKAR3 biosensor (Yapo et al., 2017).

**Figure 5.**
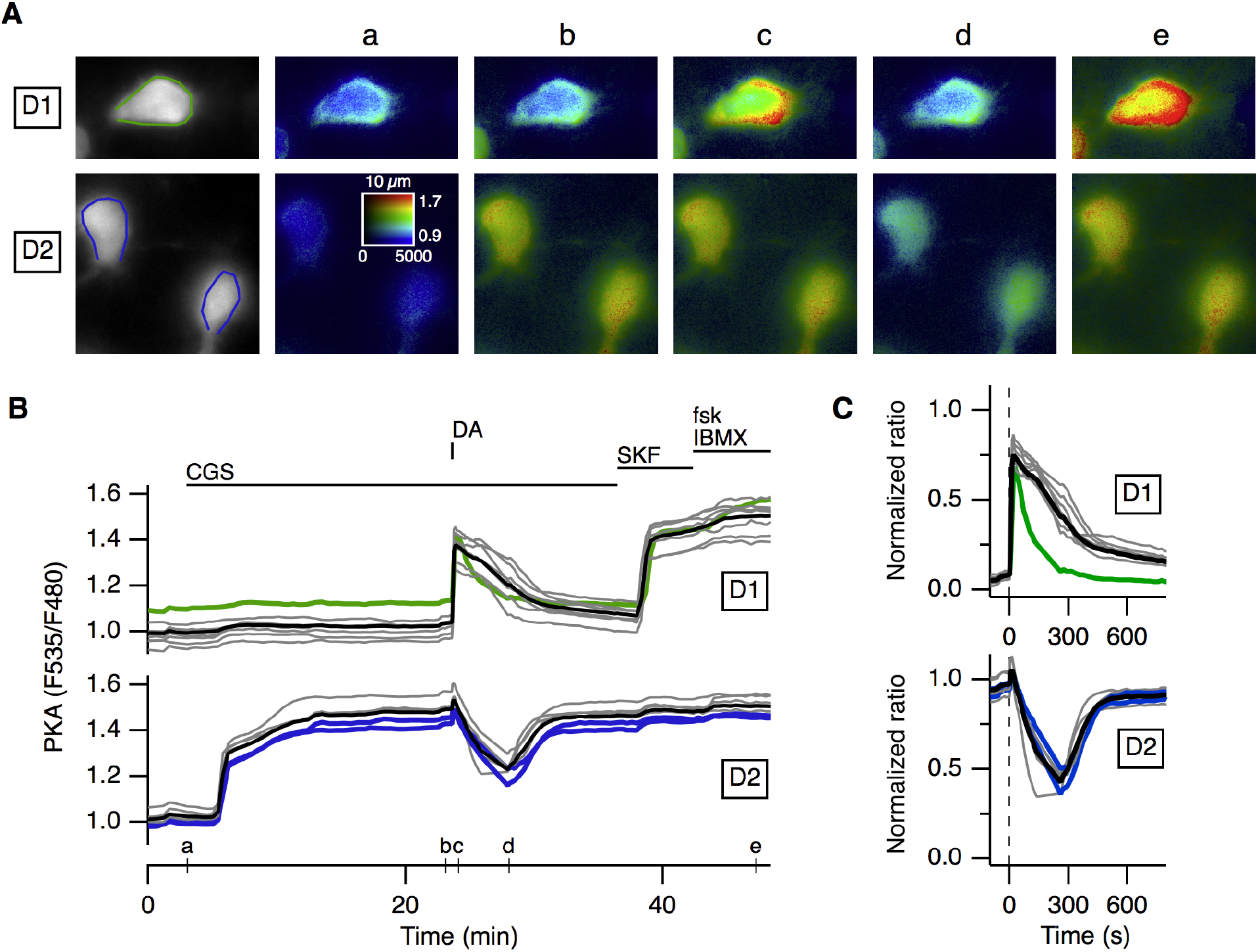
Transient dopamine induces a opposite responses at the level of PKA-dependent phosphorylation in D1 and D2 MSNs. The AKAR4 biosensor was expressed in mice striatal brain slices and imaged with wide-field microscopy. (A, left) Selected D1 and D2 MSNs on the field showing the fluorescence at the acceptor wavelength (F535), displayed in grey. The regions of interest around each MSN define the area used for measuring the ratio (F535/F480), indicative of PKA-dependent phosphorylation level. Pseudo-colour images (a-e) of the selected region corresponds to the time points indicated on the graph in B. (B) Each trace on the graph indicates the variations in the emission ratio measured on the cell body of an individual neuron. Drug application is indicated by horizontal bars. The A2a receptor agonist CGS 21680 (CGS, 1 μM) was applied to selectively activate PKA signalling in D2 cells. Dopamine (DA) release from caged NPEC-DA (3 *μ*M) by a flash of UV light triggered a transient response in D1 MSNs, and a trough in the D2 MSNs. The final application of forskolin (fsk, 13 *μ*M) and IBMX (200 *μ*M) revealed the saturating ratio level of the biosensor for each neurone. Green and blue traces indicate, respectively, the ratio variations in the D1 or D2 MSNs represented in A. (C) The traces of the individual cells in the field were normalised with respect to their maximal response to fsk and IBMX. The averages of D1 and D2 ratio traces are represented as a black line.

To test for the functional role of PDE10A, PDE2A and PDE4 at the level of PKA-dependent phosphorylation level, the same experiment was performed in the presence of TP-10 (Figure 6A) or BAY and roli (Figure 6B). This protocol consistently showed that PDE10A inhibition with TP-10 profoundly altered the responsiveness to transient dopamine (Figure 6C, D, red traces and Table 2): upon TP-10 treatment, D1 MSNs started off from a high ratio level that was not increased by dopamine nor fsk and IBMX. In this condition, transient dopamine signal was not transduced at the level of PKA-dependent phosphorylation. In D2 MSNs, transient dopamine induced little reduction in PKA signal. In contrast to TP-10, the inhibition of PDE2A and PDE4 did not significantly affect the PKA response to transient dopamine.

**Figure 6.**
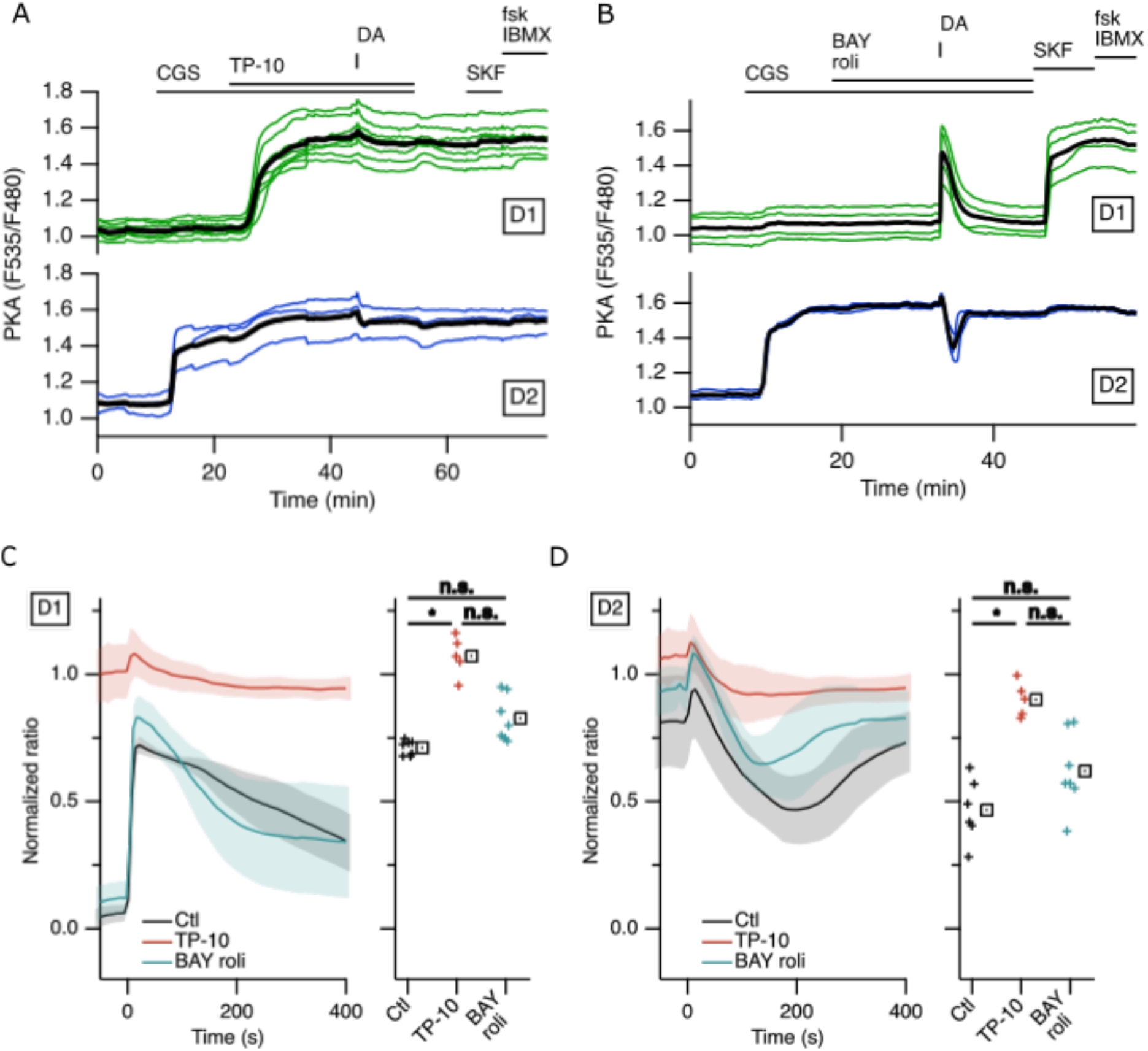
PDE10A activity is required to transduce transient dopamine at the level of PKA-dependent phosphorylation in D1 and D2 MSNs. Same experiment as in Figure 5, except that dopamine (DA) was released either in the presence of the PDE10A inhibitor TP-10 (1 *μ*M, A) or in the simultaneous presence of PDE2A inhibitor (BAY607550, BAY, 200 nM) and PDE4 inhibitor (rolipram, roli, 1 *μ*M) (B). Plots show the average of the amplitude of peak in D1 (C) and trough in D2 (D) MSNs, in control condition (ctl), in the presence of TP-10, or BAY and roli. * indicates a statistically significant difference; n.s. stands for not significantly different.

**Table 2:** Measurements corresponding to figure 6.

|  |  | Numbers |  |  | Peak/trough |
| --- | --- | --- | --- | --- | --- |
|  |  | N | n | A | ratio |
| D1 | Control | 7 | 49 | 4 | 0,711 |
|  | TP-10 | 5 | 36 | 3 | 1,072 |
|  | BAY roli | 7 | 33 | 3 | 0,827 |
| D2 | Control | 6 | 30 | 4 | 0,466 |
|  | TP-10 | 5 | 22 | 3 | 0,901 |
|  | BAY roli | 7 | 29 | 3 | 0,620 |

Switching-off cAMP production with transient dopamine is too brief to allow for a large dephosphorylation of PKA targets in D2 MSNs, as show previously (Yapo et al., 2017) and in Figure 5. Therefore, we tested with continuous bath application of dopamine which PDE activity was required to lower cAMP concentration at a level that would be sufficiently low to deactivates PKA. As with the previous protocol, the adenosine tone was mimicked with the A_2A_ agonist, CGS.

Using the cAMP biosensor Epac-S^H150^ and in the presence of TC-E, dopamine application lowered cAMP to ~3 μM in D2 MSNs (Figure 7A, E, and Table 3). In contrast, when PDE2A and PDE4 were inhibited with PF and picla, dopamine decreased cAMP concentration to a significantly lower level, close to baseline (Figure 7C, E, and Table 3).

**Figure 7.**
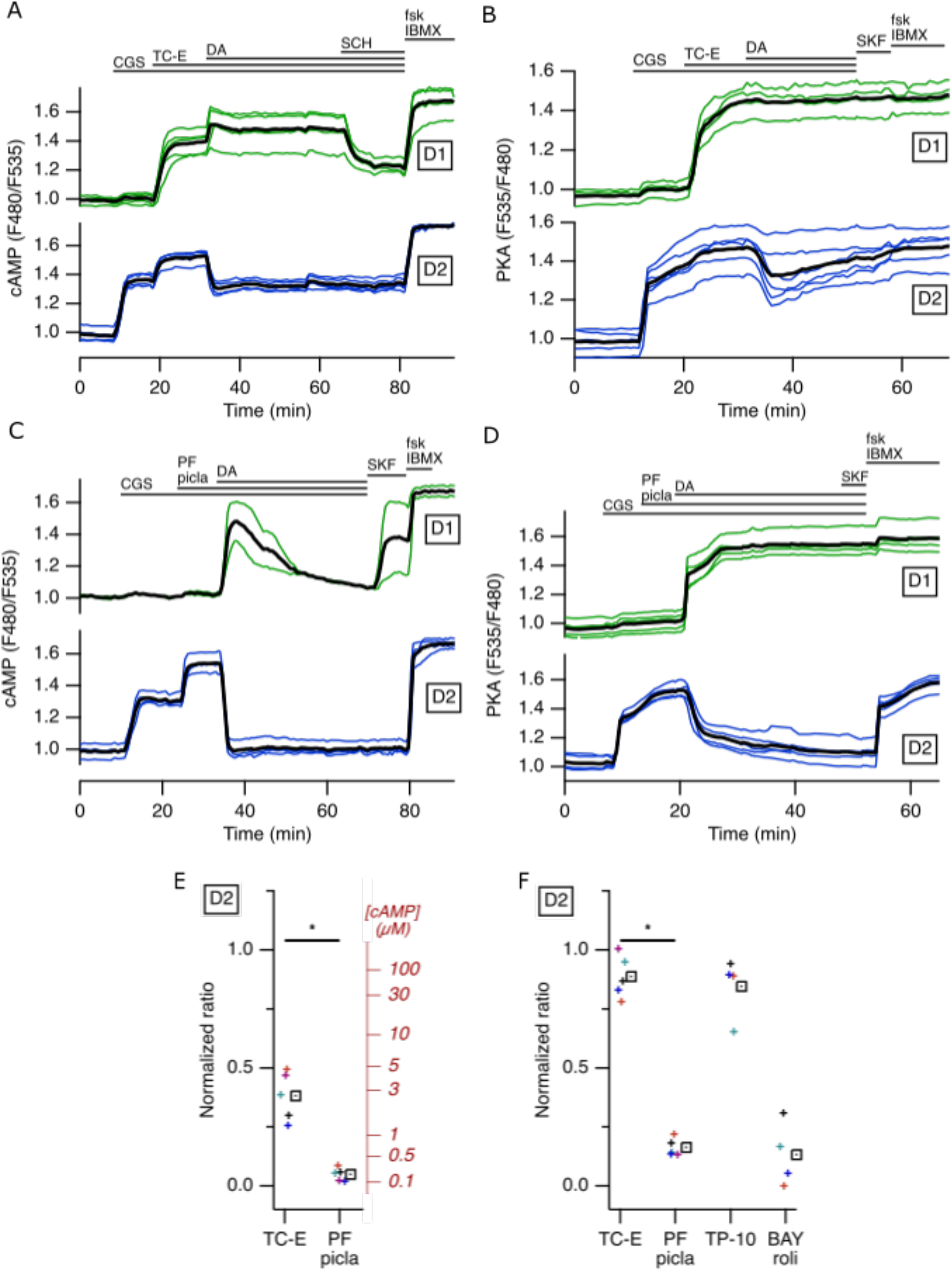
PDE10A activity is required to allow for cAMP decrease and PKA deactivation in D2 MSNs during sustained dopamine stimulus. Epac-S^H150^ (A, C) or AKAR4 (B, D) biosensors were used to monitor respectively cAMP or PKA-dependent phosphorylation changes, measured on the cell body of individual neurones. Dopamine (DA, 3 *μ*M) was applied in the bath in the presence of the PDE10A inhibitors TC-E (A, B) or in the presence of PDE2A and PDE4 inhibitors PF-05180999 (PF, 1*μ*M) and piclamilast (picla, 1*μ*M). (E) shows the average reduction in cAMP level upon PDE10A inhibition (TC-E) or PDE2 and PDE4 inhibition (PF picla). (F) shows the reduction in PKA-dependent phosphorylation level in the same condition, as well as with the other inhibitors TP-10 or BAY roli.

**Table 3:** Measurements corresponding to figure 7.

|  |  |  | Numbers |  |  | St. state |  |  |
| --- | --- | --- | --- | --- | --- | --- | --- | --- |
| | | | N | n | A | ratio | | [cAMP]<br>(μM) |
| cAMP | D2 | TC-E | 5 | 19 | 3 | 0,381 | * | 2,6419 |
|  |  | PF picla | 5 | 16 | 3 | 0,049 |  | 0,1926 |
| PKA | D2 | TC-E | 5 | 19 | 4 | 0,887 | * |  |
|  |  | PF picla | 5 | 18 | 3 | 0,163 |  |  |
|  |  | TP-10 | 4 | 9 | 6 | 0,845 |  |  |
|  |  | BAY roli | 4 | 8 | 2 | 0,133 |  |  |

The same protocol was performed using AKAR4 to monitor PKA-dependent phosphorylation level. CGS application led to a high steady-state PKA-dependent phosphorylation level in D2 MSNs. Application of the PDE10A inhibitor TC-E increased PKA-dependent phosphorylation to the maximal ratio level in both D1 and D2 MSNs. In D2 MSNs, dopamine produced a transient and moderate decrease that progressively returned to the maximal level (Figure 7B, F, blue traces). The profile of PKA-dependent phosphorylation was radically different with the inhibition of both PDE2A and PDE4 (PF and picla, respectively): sustained application of dopamine (3 μM) efficiently suppressed PKA-dependent phosphorylation level in D2 MSNs (Figure 7D, F, blue traces). The same protocol was repeated with the other inhibitors, TP-10 and BAY roli, showing similar results (Figure 7F).

This experiment demonstrates that dopamine, via D_2_ receptors, leads to the deactivation of PKA and the dephosphorylation of the kinase substrates. This effect of dopamine requires PDE10A activity, whereas PDE2A and PDE4 action appears incidental.

## Discussion

### PDE specificity on different cAMP levels

Our study reveals the specific action of PDE2A, PDE4 and PDE10A in the regulation of the cAMP/PKA signalling cascade in both D1 and D2 MSNs. Our results are quite consistent with the idea that different PDEs exhibit distinctive enzymatic properties endowing them with a specific functional role in striatal neurones (Neves-Zaph, 2017): in this review, the author highlighted that PDE2A, which exhibits a Km of several tens of micromolar, should mainly degrade the highest cAMP levels. Indeed, our experiments show that PDE2A inhibition increases the peak amplitude of cAMP responses to dopamine in D1 MSNs, as well as the A_2A_-induced steady-state cAMP level in D2 MSNs. Both levels are higher than several micromolar cAMP concentration. PDE2A however is not involved in the regulation of basal (i.e. unstimulated) cAMP since its inhibition did not affect basal cAMP level in MSNs of both types (Polito et al., 2013), whereas PDE10A inhibition did, as shown here and in our previous work (Polito et al., 2015). Our work also shows that PDE4 is functional in both D1 and D2 MSNs since its inhibition increases the cAMP responses to D_1_ or A_2A_ receptor stimulation. However, like PDE2A, PDE4 do not play a significant role in the regulation of basal cAMP. With a Km of a few micromolar, PDE4 thus regulates moderate to high cAMP levels, as suggested by computer simulations (Neves-Zaph, 2017). Concomitant inhibition of both PDE2A and PDE4 essentially exacerbated the effect of the inhibition of either one: it had no or minimal effect on basal cAMP level in D1 MSNs, but it increased the amplitude and duration of the transient cAMP response to D1 stimulation and potentiated the cAMP response to A_2A_ receptor activation in D2 MSNs. In contrast to PDE2A and PDE4, PDE10A has a sub-micromolar Km for cAMP, and is thus expected to mainly regulate the lowest cAMP levels. Indeed, we already reported that partial PDE10A inhibition in the absence of any stimulation increased cAMP from sub-micromolar to micromolar concentration, consistent with PDE10A regulating tonic cAMP production by adenylyl cyclases at sub-micromolar concentration level (Polito et al., 2015).

Surprisingly, our data show that PDE10A is also involved in the degradation of high cAMP levels since its inhibition largely prolongs dopamine D_1_ responses and increases A_2A_-induced cAMP plateau in D2 MSNs. Enzymes with low Km values would be generally expected to finetune low concentration and be saturated - and of little functional role - when the substrate concentration is high. This is true unless Vmax is high and/or the enzyme is present at high concentration. Indeed, PDE10A is expressed at very high levels and at all ages in MSNs (Seeger et al., 2003; Grauer et al., 2009; Lakics et al., 2010; Kelly et al., 2014). This also comes in good agreement with previous biochemical studies which showed that PDE10A accounted for the vast majority of cAMP hydrolysing activity in striatal extracts (Russwurm, Koesling, & Russwurm, 2015). By working on both low and high cAMP levels, PDE10A thus appears to play a predominant functional role in MSNs.

However, one should keep in mind that our measurements were performed in new born mice, and PDE expression levels change through mouse and human lifespan (Kelly et al., 2014; Hegde et al., 2016; Farmer et al., 2020). Since PDE10A remains the dominant form of PDE in the adult striatum, it is quite plausible that its key signalling function is maintained in adulthood.

### Signal integration at the level of PKA-dependent phosphorylation

PKA, the primary target of cAMP in MSNs, is activated by sub-micromolar cAMP levels (Dostmann & Taylor, 1991). PDE10A is the only striatal PDE working on sub-micromolar cAMP levels, and our data confirm that its activity is required for proper integration of cAMP signals at the PKA level. In D1 MSNs, transient dopamine triggers a transient increase in PKA-dependent phosphorylation level, and such transient signals are essentially abolished when PDE10A is inhibited: baseline cAMP level increases above PKA activation threshold, and any further stimulation of cAMP production, for example through D_1_ receptors, has no significant effect on a continuously elevated PKA signal. This effect is expected to be strongly potentiated by the positive feed-forward control mediated by DARPP-32 (Girault & Greengard, 2004), which prevents phosphatase action. Such mechanism promotes an all-or-none responsiveness in D1 MSNs, as has been described in the nucleus of MSNs (Yapo et al., 2018). This locks D1 MSNs in a highly phosphorylated state, likely associated with elevated activity in the direct pathway.

D2 MSNs are intrinsically not geared to respond to transient dopamine signals, since it takes time to transduce the D_2_-induced decrease in cAMP into a dephosphorylation of PKA substrates. This has been reported previously (Yapo et al., 2017) and illustrated here in Figure 5. Modelling analysis suggested that D2 MSNs might detect a transient absence of dopamine, in a context of tonic A_2A_ receptor stimulation (Yapo et al., 2017). This mechanism requires that tonic dopamine maintains a low PKA-dependent phosphorylation level even in the presence of adenosine tone. Figure 7 shows that this cannot be obtained if PDE10A is inhibited: cAMP cannot be maintained below micromolar level, and PKA-dependent phosphorylation level remains high.

Besides PDE10A activity, which appears as the centrepiece of PKA signalling in MSNs, PDE2A and PDE4 display ancillary effects that may nonetheless bear important functional significance. First, PDE2A and PDE4 affect the amplitude of cAMP peaks in D1 MSNs. It was previously shown that PDE1B also regulates the peak cAMP response to dopamine, an effect that translated into changes in synaptic plasticity (Betolngar et al., 2019). Similarly, PDE2A activation through the NO/cGMP signalling axis has also been reported to modulate transient PKA responses (Polito et al., 2013). Fine-tuning of peak PKA responses through PDE2A and PDE4 may therefore be of importance in the outcome of phasic dopamine signals in the striatum.

This work also demonstrates that PDE2A and PDE4 regulate steady-state micromolar cAMP levels. This action may have important functional effects by determining thresholds for switching from low to high PKA-dependent phosphorylation level, as shown in the D_2_/A_2A_ interaction (Yapo et al., 2017). Further experimental and modelling work would be necessary to precise these effects. Finally, although this has not been characterised precisely in this work, PDE2A and PDE4 contribute to cAMP degradation after activation of D_2_ receptors, and their inhibition slows down the return towards baseline. PDE2A and PDE4 modulation is therefore expected to induce changes in the relative timing of signalling events, which is a critical aspect of striatal function.

### PDEs as therapeutic targets

As indicated in the introduction, inhibiting PDE10A seemed an interesting approach by its selectivity for striatal neurones. Our data show that this strategy exposes to a delicate situation in which the therapeutic window can be very narrow, between an expected beneficial effect, such as boosting the activity of D2 MSNs, and unwanted side effects which may arise from switching a variable fraction of D1 MSNs into a highly phosphorylated state. Indeed, with MSNs responding in a very non-linear manner, it can be expected that, above a certain threshold of PDE10A inhibition, some neurones may abruptly switch to a high PKA-dependent phosphorylation state for a prolonged duration. This may underlie the unwanted side-effects observed in patients treated with high doses of PDE10A inhibitors, which led to abandoning this approach in several drug development programs (Baillie, Tejeda, & Kelly, 2019). On the other hand, regulating the other PDEs such as PDE2A and PDE4 may provide a more subtle way to adjust the cAMP response and therefore tune PKA responses. For example, behavioral studies indicate that PDE2A inhibitors may be of interest for treating cognitive impairments including those associated with schizophrenia (Boess et al., 2004; Redrobe et al., 2014; Redrobe et al., 2015), while changes in striatal PDE2A expression level were correlated with psychiatric diseases (Farmer et al., 2020).

## Conclusion

We observed that PDE2A, PDE4 and PDE10A play distinct roles in the regulation of cAMP/ PKA responses to dopamine in D1 and D2 MSNs. We report that PDE10A is required for degrading low cAMP levels which is necessary to return PKA substrates to the dephosphorylated state, while the modulation of PDE2A and PDE4 may provide a more subtle control.

## Author contribution

Conceptualization, L.C. and P.V.; Data curation, E.M., L.C. and P.V.; Formal analysis, P.V.; Funding acquisition, L.C. and P.V.; Investigation, E.M. and S.B.; Methodology, E.M. and P.V.; Project administration, P.V.; Resources, E.M., D.B., L.C. and P.V.; Software, P.V.; Supervision, P.V.; Validation, E.M. and P.V.; Visualization, E.M. and P.V.; Writing - original draft preparation, E.M., L.C. and P.V.; Writing - review and editing, E.M., L.C. and P.V. All authors have read and agreed to the published version of the manuscript.

## Funding

This research was funded by “Investissements d’Avenir” program managed by the ANR under reference ANR-11-IDEX-0004-02, and called “Bio-Psy Labex”. E.M. benefited from a PhD fellowship from this institution.

## Conflict of Interest

The authors declare no conflict of interest. The funders had no role in the design of the study; in the collection, analyses, or interpretation of data, in the writing of the manuscript, or in the decision to publish the results.

## Ethics

This publications follows the guidelines from the Committee on Publication Ethics (COPE).

## Data availability

The data that support the findings of this study are available from the corresponding author upon reasonable request.

## Acknowledgments

The team is a member of the Bio-Psy Labex. We thank Cédric Yapo, Manon Dobrigna and Ilyes Nedjar for helpful contributions to the experimental work. We thank Mohamed Doulazmi for helpful discussion on the statistical analysis.

## Notes

### Competing Interest Statement

The authors have declared no competing interest.

## References

Baillie, G. S., Tejeda, G. S., & Kelly, M. P. (2019). Therapeutic targeting of 3’,5’-cyclic nucleotide phosphodiesterases: inhibition and beyond. Nat Rev Drug Discov, 18(10), 770–796.

Bender, A. T., & Beavo, J. A. (2006). Cyclic nucleotide phosphodiesterases: molecular regulation to clinical use. Pharmacol Rev, 58(3), 488–520.

Bertran-Gonzalez, J., Hervé, D., Girault, J. A., & Valjent, E. (2010). What is the Degree of Segregation between Striatonigral and Striatopallidal Projections. Front Neuroanat, 4, 136.

Betolngar, D. B., Mota, É., Fabritius, A., Nielsen, J., Hougaard, C., Christoffersen, C. T., Yang, J., Kehler, J., Griesbeck, O., Castro, L. R. V., & Vincent, P. (2019). Phosphodiesterase 1 Bridges Glutamate Inputs with NO- and Dopamine-Induced Cyclic Nucleotide Signals in the Striatum. Cereb Cortex, 29(12), 5022–5036.

Boess, F. G., Hendrix, M., van der Staay, F. J., Erb, C., Schreiber, R., van Staveren, W., de Vente, J., Prickaerts, J., Blokland, A., & Koenig, G. (2004). Inhibition of phosphodiesterase 2 increases neuronal cGMP, synaptic plasticity and memory performance. Neuropharmacology, 47(7), 1081–1092.

Cadd, G., & McKnight, G. S. (1989). Distinct patterns of cAMP-dependent protein kinase gene expression in mouse brain. Neuron, 3(1), 71–79.

Cerovic, M., d’Isa, R., Tonini, R., & Brambilla, R. (2013). Molecular and cellular mechanisms of dopamine-mediated behavioral plasticity in the striatum. Neurobiol Learn Mem, 105, 63–80.

Chappie, T., Humphrey, J., Menniti, F., & Schmidt, C. (2009). PDE10A inhibitors: an assessment of the current CNS drug discovery landscape. Curr Opin Drug Discov Devel, 12(4), 458–467.

Cherry, J. A., & Davis, R. L. (1999). Cyclic AMP phosphodiesterases are localized in regions of the mouse brain associated with reinforcement, movement, and affect. J Comp Neurol, 407(2), 287–301.

Coskran, T. M., Morton, D., Menniti, F. S., Adamowicz, W. O., Kleiman, R. J., Ryan, A. M., Strick, C. A., Schmidt, C. J., & Stephenson, D. T. (2006). Immunohistochemical localization of phosphodiesterase 10A in multiple mammalian species. J Histochem Cytochem, 54(11), 1205–1213.

Curtis, M. J., Alexander, S., Cirino, G., Docherty, J. R., George, C. H., Giembycz, M. A., Hoyer, D., Insel, P. A., Izzo, A. A., Ji, Y., MacEwan, D. J., Sobey, C. G., Stanford, S. C., Teixeira, M. M., Wonnacott, S., & Ahluwalia, A. (2018). Experimental design and analysis and their reporting II: updated and simplified guidance for authors and peer reviewers. Br J Pharmacol, 175(7), 987–993.

Day, J. J., Roitman, M. F., Wightman, R. M., & Carelli, R. M. (2007). Associative learning mediates dynamic shifts in dopamine signaling in the nucleus accumbens. Nat Neurosci, 10(8), 1020–1028.

Depry, C., Allen, M. D., & Zhang, J. (2011). Visualization of PKA activity in plasma membrane microdomains. Mol Biosyst, 7(1), 52–58.

Dostmann, W. R., & Taylor, S. S. (1991). Identifying the molecular switches that determine whether (Rp)-cAMPS functions as an antagonist or an agonist in the activation of cAMP-dependent protein kinase I. Biochemistry, 30(35), 8710–8716.

Ehrengruber, M. U., Lundstrom, K., Schweitzer, C., Heuss, C., Schlesinger, S., & Gähwiler, B. H. (1999). Recombinant Semliki Forest virus and Sindbis virus efficiently infect neurons in hippocampal slice cultures. Proc Natl Acad Sci U S A, 96(12), 7041–7046.

Farmer, R., Burbano, S. D., Patel, N. S., Sarmiento, A., Smith, A. J., & Kelly, M. P. (2020). Phosphodiesterases PDE2A and PDE10A both change mRNA expression in the human brain with age, but only PDE2A changes in a region-specific manner with psychiatric disease. Cell Signal, 70, 109592.

Fink, J. S., Weaver, D. R., Rivkees, S. A., Peterfreund, R. A., Pollack, A. E., Adler, E. M., & Reppert, S. M. (1992). Molecular cloning of the rat A2 adenosine receptor: selective co-expression with D2 dopamine receptors in rat striatum. Brain Res Mol Brain Res, 14(3), 186–195.

Girault, J. A., & Greengard, P. (2004). The neurobiology of dopamine signaling. Arch Neurol, 61(5), 641–644.

Gonon, F. (1997). Prolonged and extrasynaptic excitatory action of dopamine mediated by D1 receptors in the rat striatum in vivo. J Neurosci, 17(15), 5972–5978.

Grauer, S. M., Pulito, V. L., Navarra, R. L., Kelly, M., Kelley, C., Graf, R., Langen, B., Logue, S., Brennan, J., Jiang, L., Charych, E., Egerland, U., Liu, F., Marquis, K. L., Malamas, M., Hage, T., Comery, T. A., & Brandon, N. J. (2009). PDE10A Inhibitor Activity in Preclinical Models of the Positive, Cognitive and Negative Symptoms of Schizophrenia. J Pharmacol Exp Ther.

Grynkiewicz, G., Poenie, M., & Tsien, R. Y. (1985). A new generation of Ca2+ indicators with greatly improved fluorescence properties. J Biol Chem, 260(6), 3440–3450.

Harada, A., Kaushal, N., Suzuki, K., Nakatani, A., Bobkov, K., Vekich, J. A., Doyle, J. P., & Kimura, H. (2020). Balanced Activation of Striatal Output Pathways by Faster Off-Rate PDE10A Inhibitors Elicits Not Only Antipsychotic-Like Effects But Also Procognitive Effects in Rodents. Int J Neuropsychopharmacol, 23(2), 96–107.

Hegde, S., Capell, W. R., Ibrahim, B. A., Klett, J., Patel, N. S., Sougiannis, A. T., & Kelly, M. P. (2016). Phosphodiesterase 11A (PDE11A), Enriched in Ventral Hippocampus Neurons, is Required for Consolidation of Social but not Nonsocial Memories in Mice. Neuropsychopharmacology.

Jäger, R., Schwede, F., Genieser, H. G., Koesling, D., & Russwurm, M. (2010). Activation of PDE2 and PDE5 by specific GAF ligands: delayed activation of PDE5. Br J Pharmacol, 161(7), 1645–1660.

Kehler, J., & Nielsen, J. (2011). PDE10A inhibitors: novel therapeutic drugs for schizophrenia. Curr Pharm Des, 17(2), 137–150.

Kehler, J., Ritzén, A., & Greve, D. R. P. Q. (2007). The potential therapeutic use of phosphodiesterase 10 inhibitors. Expert Opin. Ther. Patents, 17(2), 147–158.

Kelly, M. P., Adamowicz, W., Bove, S., Hartman, A. J., Mariga, A., Pathak, G., Reinhart, V., Romegialli, A., & Kleiman, R. J. (2014). Select 3’,5’-cyclic nucleotide phosphodiesterases exhibit altered expression in the aged rodent brain. Cell Signal, 26(2), 383–397.

Klaus, A., Alves da Silva, J., & Costa, R. M. (2019). What, If, and When to Move: Basal Ganglia Circuits and Self-Paced Action Initiation. Annu Rev Neurosci, 42, 459–483.

Kreitzer, A. C., & Malenka, R. C. (2008). Striatal plasticity and basal ganglia circuit function. Neuron, 60(4), 543–554.

Lakics, V., Karran, E. H., & Boess, F. G. (2010). Quantitative comparison of phosphodiesterase mRNA distribution in human brain and peripheral tissues. Neuropharmacology, 59(6), 367–374.

Martinez, S. E., Wu, A. Y., Glavas, N. A., Tang, X. B., Turley, S., Hol, W. G., & Beavo, J. A. (2002). The two GAF domains in phosphodiesterase 2A have distinct roles in dimerization and in cGMP binding. Proc Natl Acad Sci U S A, 99(20), 13260–13265.

Martins, T. J., Mumby, M. C., & Beavo, J. A. (1982). Purification and characterization of a cyclic GMP-stimulated cyclic nucleotide phosphodiesterase from bovine tissues. J Biol Chem, 257(4), 1973–1979.

Menniti, F. S., Chappie, T. A., Humphrey, J. M., & Schmidt, C. J. (2007). Phosphodiesterase 10A inhibitors: a novel approach to the treatment of the symptoms of schizophrenia. Curr Opin Investig Drugs, 8(1), 54–59.

Mika, D., Bobin, P., Lindner, M., Boet, A., Hodzic, A., Lefebvre, F., Lechène, P., Sadoune, M., Samuel, J. L., Algalarrondo, V., Rucker-Martin, C., Lambert, V., Fischmeister, R., Vandecasteele, G., & Leroy, J. (2019). Synergic PDE3 and PDE4 control intracellular cAMP and cardiac excitation-contraction coupling in a porcine model. J Mol Cell Cardiol, 133, 57–66.

Neves-Zaph, S. R. (2017). Phosphodiesterase Diversity and Signal Processing Within cAMP Signaling Networks. Adv Neurobiol, 17, 3–14.

Nishi, A., Kuroiwa, M., Miller, D. B., O’Callaghan, J. P., Bateup, H. S., Shuto, T., Sotogaku, N., Fukuda, T., Heintz, N., Greengard, P., & Snyder, G. L. (2008). Distinct roles of PDE4 and PDE10A in the regulation of cAMP/PKA signaling in the striatum. J Neurosci, 28(42), 10460–10471.

Nishi, A., & Snyder, G. L. (2010). Advanced research on dopamine signaling to develop drugs for the treatment of mental disorders: biochemical and behavioral profiles of phosphodiesterase inhibition in dopaminergic neurotransmission. J Pharmacol Sci, 114(1), 6–16.

Paes, D., Xie, K., Wheeler, D. G., Zook, D., Prickaerts, J., & Peters, M. (2021). Inhibition of PDE2 and PDE4 synergistically improves memory consolidation processes. Neuropharmacology, 184, 108414.

Perez-Torres, S., Miro, X., Palacios, J. M., Cortes, R., Puigdomenech, P., & Mengod, G. (2000). Phosphodiesterase type 4 isozymes expression in human brain examined by in situ hybridization histochemistry and[3H]rolipram binding autoradiography. Comparison with monkey and rat brain. J Chem Neuroanat, 20(3-4), 349–374.

Polito, M., Guiot, E., Gangarossa, G., Longueville, S., Doulazmi, M., Valjent, E., Hervé, D., Girault, J. A., Paupardin-Tritsch, D., Castro, L. R., & Vincent, P. (2015). Selective Effects of PDE10A Inhibitors on Striatopallidal Neurons Require Phosphatase Inhibition by DARPP-32. eNeuro, 2(4), 1–15.

Polito, M., Klarenbeek, J., Jalink, K., Paupardin-Tritsch, D., Vincent, P., & Castro, L. R. (2013). The NO/ cGMP pathway inhibits transient cAMP signals through the activation of PDE2 in striatal neurons. Front Cell Neurosci, 7, 211.

Polli, J. W., & Kincaid, R. L. (1994). Expression of a calmodulin-dependent phosphodiesterase isoform (PDE1B1) correlates with brain regions having extensive dopaminergic innervation. J Neurosci, 14(3 Pt 1), 1251–1261.

Poppe, H., Rybalkin, S. D., Rehmann, H., Hinds, T. R., Tang, X. B., Christensen, A. E., Schwede, F., Genieser, H. G., Bos, J. L., Doskeland, S. O., Beavo, J. A., & Butt, E. (2008). Cyclic nucleotide analogs as probes of signaling pathways. Nat Methods, 5(4), 277–278.

Redrobe, J. P., Jørgensen, M., Christoffersen, C. T., Montezinho, L. P., Bastlund, J. F., Carnerup, M., Bundgaard, C., Lerdrup, L., & Plath, N. (2014). In vitro and in vivo characterisation of Lu AF64280, a novel, brain penetrant phosphodiesterase (PDE) 2A inhibitor: potential relevance to cognitive deficits in schizophrenia. Psychopharmacology (Berl), 231(16), 3151–3167.

Redrobe, J. P., Rasmussen, L. K., Christoffersen, C. T., Bundgaard, C., & Jorgensen, M. (2015). Characterisation of Lu AF33241: A novel, brain-penetrant, dual inhibitor of phosphodiesterase (PDE) 2A and PDE10A. Eur J Pharmacol.

Russwurm, C., Koesling, D., & Russwurm, M. (2015). Phosphodiesterase 10A Is Tethered to a Synaptic Signaling Complex in Striatum. J Biol Chem, 290(19), 11936–11947.

Schülke, J. P., & Brandon, N. J. (2017). Current Understanding of PDE10A in the Modulation of Basal Ganglia Circuitry. Adv Neurobiol, 17, 15–43.

Schultz, W., Dayan, P., & Montague, P. R. (1997). A neural substrate of prediction and reward. Science, 275(5306), 1593–1599.

Seeger, T. F., Bartlett, B., Coskran, T. M., Culp, J. S., James, L. C., Krull, D. L., Lanfear, J., Ryan, A. M., Schmidt, C. J., Strick, C. A., Varghese, A. H., Williams, R. D., Wylie, P. G., & Menniti, F. S. (2003). Immunohistochemical localization of PDE10A in the rat brain. Brain Res, 985(2), 113–126.

Stephenson, D. T., Coskran, T. M., Kelly, M. P., Kleiman, R. J., Morton, D., O’Neill, S. M., Schmidt, C. J., Weinberg, R. J., & Menniti, F. S. (2012). The distribution of phosphodiesterase 2A in the rat brain. Neuroscience, 226, 145–155.

Stephenson, D. T., Coskran, T. M., Wilhelms, M. B., Adamowicz, W. O., O’Donnell, M. M., Muravnick, K. B., Menniti, F. S., Kleiman, R. J., & Morton, D. (2009). Immunohistochemical localization of phosphodiesterase 2A in multiple mammalian species. J Histochem Cytochem, 57(10), 933–949.

Sudo, T., Tachibana, K., Toga, K., Tochizawa, S., Inoue, Y., Kimura, Y., & Hidaka, H. (2000). Potent effects of novel anti-platelet aggregatory cilostamide analogues on recombinant cyclic nucleotide phosphodiesterase isozyme activity. Biochem Pharmacol, 59(4), 347–356.

Surmeier, D. J., Ding, J., Day, M., Wang, Z., & Shen, W. (2007). D1 and D2 dopamine-receptor modulation of striatal glutamatergic signaling in striatal medium spiny neurons. Trends Neurosci, 30(5), 228–235.

Threlfell, S., Sammut, S., Menniti, F. S., Schmidt, C. J., & West, A. R. (2009). Inhibition of Phosphodiesterase 10A Increases the Responsiveness of Striatal Projection Neurons to Cortical Stimulation. J Pharmacol Exp Ther, 328(3), 785–795.

Valjent, E., Bertran-Gonzalez, J., Herve, D., Fisone, G., & Girault, J. A. (2009). Looking BAC at striatal signaling: cell-specific analysis in new transgenic mice. Trends Neurosci, 32(10), 538–547.

Wang, H., Liu, Y., Hou, J., Zheng, M., Robinson, H., & Ke, H. (2007). Structural insight into substrate specificity of phosphodiesterase 10. Proc Natl Acad Sci U S A, 104(14), 5782–5787.

Yan, C., Bentley, J. K., Sonnenburg, W. K., & Beavo, J. A. (1994). Differential expression of the 61 kDa and 63 kDa calmodulin-dependent phosphodiesterases in the mouse brain. J Neurosci, 14(3 Pt 1), 973–84.

Yapo, C., Nair, A. G., Clement, L., Castro, L. R., Hellgren Kotaleski, J., & Vincent, P. (2017). Detection of phasic dopamine by D1 and D2 striatal medium spiny neurons. J Physiol, 595(24), 7451–7475.

Yapo, C., Nair, A. G., Hellgren Kotaleski, J., Vincent, P., & Castro, L. R. V. (2018). Switch-like PKA responses in the nucleus of striatal neurons. J Cell Sci, 131(14), jcs.216556.

